# The role of contextual similarity on brain activity patterns for associated events

**DOI:** 10.64898/2026.10.19.760268

**Authors:** L. Gurguryan, J. M. Olivier, H. R Dimsdale-Zucker

## Abstract

Our everyday lives are characterized by repeated contexts, such as returning to work in the same office every day. Although the physical office may be identical, other features vary— e.g., time of day, light conditions, one’s internal state—leading to differing degrees of similarity, even among putatively identical contexts. Alternatively, contexts can change abruptly, such as leaving the office to go home for dinner. Despite the prevalence of various levels of contextual overlap in everyday life, prior experimental work has largely ignored their influence on related experiences. To address this gap, we took a novel approach in which participants viewed real-world audiovisual stimuli (commercials [events], TV shows [contexts]) while undergoing functional magnetic resonance imaging scans (fMRI) to examine whether the degree of contextual overlap influenced brain activity patterns for related events. Representational similarity analyses revealed sensitivity to global context—greater similarity for commercials seen within the same versus different show—in ventromedial prefrontal cortex, posterior medial cortex, perirhinal cortex, and parahippocampal cortex. Additionally, parahippocampal cortex was uniquely sensitive to local context—greater similarity for commercials seen in the same versus different episode of the same show—and showed reduced activity to repetition of event-specific information, suggesting representation of event-level conceptual information. Thus, even with no explicit demand to retrieve contextual associations, event representations changed based on the degree of contextual similarity, suggesting that contextual information is an inherent part of how experiences are represented.

## Introduction

Events contain rich, multidimensional contexts which are encoded and represented in memory (Godden & Baddeley, 1975; Hockley, 2008; Smith & Manzano, 2010; Tulving & Thomson, 1973). Theories of episodic memory suggest that these context representations— which can carry information about related temporal, spatial, and even conceptual associations (Copara et al., 2014; Diana et al., 2012)—are reinstated during the retrieval of events (Herweg et al., 2020; Howard & Kahana, 2002; Polyn et al., 2009). However, it remains unclear how the brain represents contexts between events when the degree of contextual overlap varies and whether this occurs when there is no explicit memory demand for contextual information.

Retrieved-context frameworks of episodic memory propose that representations at retrieval carry associated information about the position, order, spatial and semantic associations of the memoranda that were initially instantiated at the time of encoding (Howard & Kahana, 2002). This reinstated contextual information can serve as a cue, leading to heightened retrieval of contextually similar items (Lohnas et al., 2015; Polyn et al., 2009). Additionally, memory performance is better when encoding and retrieval contexts overlap (Godden & Baddeley, 1975; Hockley, 2008; Smith, 1979; Smith & Manzano, 2010); however, see Murre, 2021; Wälti et al., 2019), or when context is reinstated explicitly (e.g., with cues; Kraflca & Penrod, 1985; Sekeres et al., 2016).

Brain imaging studies have shown that the strength of neural reinstatement—the extent to which neural activity patterns during retrieval resemble encoding-related patterns (Danker & Anderson, 2010; Rissman & Wagner, 2012; Rugg et al., 2015)—is positively associated with memory performance, and therefore thought to reflect the quantity and quality of information retrieved from memory (Gordon et al., 2014; Johnson et al., 2009; Olivier et al., 2025; Thakral et al., 2015; Trelle et al., 2020). When contextual information is explicitly manipulated, neural activity patterns associated with the encoding context can also be reinstated at retrieval (Bornstein & Norman, 2017; Bramão et al., 2017; Chan et al., 2017; Essoe et al., 2022; Lohnas et al., 2023; Masís-Obando et al., 2026) and the degree of contextual reinstatement can predict likelihood of correct recall (Manning et al., 2011).

While these findings suggest that context-associated neural activity patterns can be reliably identified, it is unclear how the brain represents and distinguishes between similar contexts. For example, if a member of a search committee is interviewing candidates, she may interview multiple different individuals in the same location (e.g., her office) but at different times (e.g., at 10 a.m. and 1 p.m., but both on Tuesday). For each of these interviews, although the spatial context (her office) remains stable, both the temporal (time of meeting) and conceptual (candidate identity, topics discussed) contexts have drifted (for a theoretical discussion of "drifting" versus "shifting" contexts, see DuBrow et al., 2017). If she had a third interview in her office with yet another candidate but at a temporally distinct context (e.g., the next day) or even in a distinct spatial and temporal context (e.g., at dinner later that evening), the level of contextual similarity between these related events would vary even more. That is, candidates that were interviewed in the same spatial location at temporally near slots share greater contextual similarity with candidates interviewed in distinct spatial locations. How then might the brain represent these experiences that vary in their degree of contextual overlap?

Libby et al. (2019) explored neural representations for memories either when the same items were present in different contexts (akin to interviewing the same candidate in your office and also at a group dinner) or when different items were present across overlapping contexts (akin to interviewing multiple candidates in your office) and found that the hippocampus represented events more similarly when they shared both item and context information, relative to events that shared only item or context information. Similarly, Chanales et al. (2017) observed that the hippocampus discriminated between overlapping contexts—routes in a naturalistic route-learning paradigm—while representing non-overlapping contexts as more similar. Additional studies with manipulations of context have observed similar findings for the hippocampus in both humans and non-human animals, supporting the idea that the hippocampus either separates similar contexts into orthogonal representations (Dimsdale-Zucker et al., 2018, 2022) or categorizes them together (Alme et al., 2014; I. Lee et al., 2004; Neunuebel et al., 2013; Wills et al., 2005). However, the majority of prior work in this domain has focused on how brain activity represents events when contextual information is presented concurrently with studied items and when there is an explicit demand to retrieve this earlier experience. No other work, to our knowledge, has investigated how varying degrees of context overlap influence item representations. That is, how might the brain represent the 10 a.m. and 1 p.m. interview candidates where some aspects of contextual information are shared (your office) but where other contextual features have changed (the meeting time, surrounding events, your mental state), and how might this differ from representations where there has been a clearer change to a distinct context (meetings in your office vs. at the restaurant).

To address the question of whether varying degrees of contextual similarity have a differential impact on brain activity patterns, we embedded commercials (events) within TV show episodes (contexts) during a functional magnetic resonance imaging (fMRI) session (**Figure 1**). Our experimental design allowed us to examine neural representations of events that share different levels of contextual overlap: commercials presented within the *same context* (same TV show, same episode), *similar context* (same TV show, different episode), or *different context* (different TV shows). We separately measured brain activity for these commercials prior to and following context exposure. Importantly, at neither the pre-context nor post-context association was there any explicit memory-related demand or task placed on these items. That is, we measured the underlying neural representation of these items while participants completed a task with no explicit memory demands. Then in our analyses, we considered whether being affiliated with the same, similar, or different contexts changed representations for these items.

**Figure 1.**
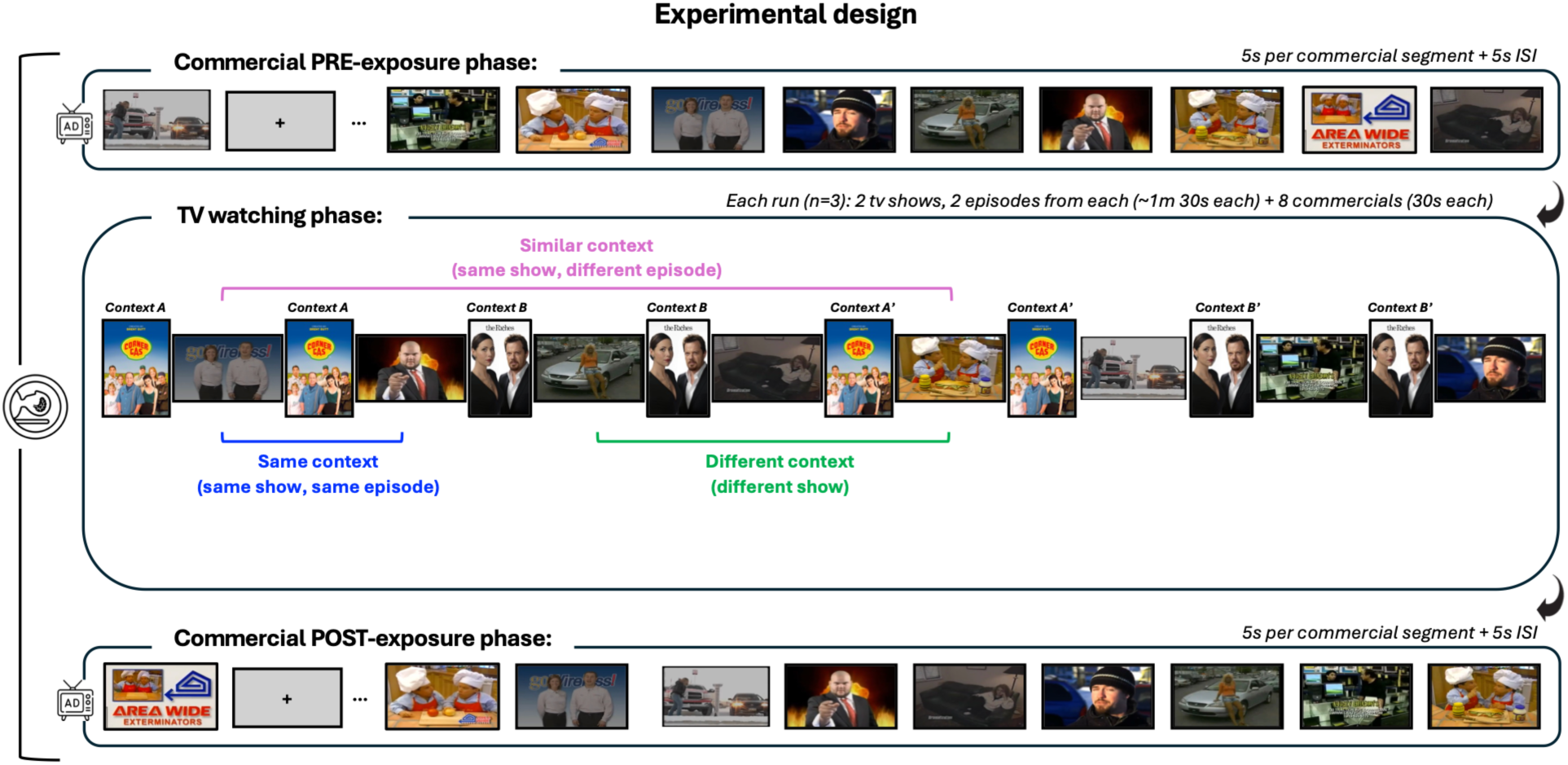
Experimental design. *Commercial PRE-exposure phase*: First, participants completed a pre-exposure run in which they viewed three 5-second segments extracted from each of 24 commercials, for a total of 72 segments, presented in a uniquely randomized order for each participant. Because the three segments from each commercial were presented in a randomized order, they did not necessarily follow their original chronological sequence as in the full version of the commercial. Participants completed a 1-back task in which they pressed a button when they believed two consecutive segments were from the same commercial. *TV watching phase:* During each of the three TV-watching runs, participants watched two episodes from two of the six possible TV shows. Complete versions of the commercials were interspersed throughout the episodes at designated commercial timepoints. These complete commercials included the three 5-second segments presented during the pre-exposure run, along with all other portions of the original commercials, presented in their original sequence. As a cover task, participants were told to pretend they had been hired from a marketing firm and to judge whether each commercial was a good fit for the TV show they had just been watching. They were told they could use content and/or timing of placement for making these judgments, but to apply a consistent judgment criterion for all commercials. *Commercial POST-exposure run*: Lastly, participants completed a post-exposure run consisting of the same 1-back task as the pre-exposure run, with the same commercial segments as the pre-exposure run presented in a different randomized order.

We hypothesized that fMRI-associated activity patterns in the ventromedial prefrontal cortex (vmPFC) and posterior medial cortex (PMC)—regions associated with forming situation models that link across experiences (Kragel et al., 2021; Ranganath & Ritchey, 2012; Ritchey et al., 2015; Ritchey & Cooper, 2020)—would be particularly sensitive to global context. That is, we hypothesized that these regions would try to represent commonalities for commercials studied within the same TV show, regardless of episode, and distinguish them from commercials affiliated with different TV shows (same context [same episode and same TV show] > different context [different TV shows]). In regions like the hippocampus that have been shown to represent individual experiences (Chanales et al., 2017; Davachi, 2006; Diana et al., 2007; Dimsdale-Zucker et al., 2022; Libby et al., 2019; Ranganath, 2010), we hypothesized that brain activity patterns would be sensitive to local context. That is, we expected that the hippocampus would represent commercials affiliated with the same episode of the same TV show as more similar to one another than if only show identity was shared (same context [same episode and same TV show] > similar context [different episodes of the same TV show]). We expected activity patterns in the parahippocampal cortex (PHC) to be sensitive to the degree to which context overlapped (same context > similar context and similar context > different context) because it is a key region in the posterior medial network which is thought to represent the contextual details of events that are combined to form situation models (Diana et al., 2013; Hannula et al., 2013; Ranganath & Ritchey, 2012; Ritchey et al., 2015; Ritchey & Cooper, 2020; Turk-Browne et al., 2012). By contrast, given the role of anterior temporal regions in item-level representations (Davachi, 2006; Diana et al., 2012; Hannula et al., 2013; Ranganath, 2010; Ranganath & Ritchey, 2012; Ritchey et al., 2015; Ritchey & Cooper, 2020; Staresina et al., 2011) we predicted that the perirhinal cortex (PRC) would be insensitive to either local or global contextual associations.

## Results

### Behavioral results

Participants correctly indicated which TV show the commercial was paired with for 45.2% of all commercials (*M* = .452, *SD* = .162). Of note, given that participants selected the associated TV show from six alternatives, chance performance was 16.7% (1/6). To parallel our RSA analyses, we also considered pair-wise source memory accuracy based on the degree of contextual overlap (same context [same TV show, same episode], similar context [same TV show, different episodes], different context [different TV show]). The mean pair-level source memory accuracy was 22.9% (*SD* = .173) for same context pairs, 21.0% (*SD* = .177) for similar context pairs, and 22.1% (*SD* = .169) for different context pairs.

### fMRI results

#### Representational similarity analyses

In order to understand how affiliation with a TV show influenced representational patterns for the associated commercials, we compared pattern similarity for local context (same TV show, same episode vs. same TV show, different episodes) and global context (same TV show, same episode vs. different TV show) trial pairs.

In ventromedial prefrontal cortex and posterior medial cortex, there was a significant overall effect of condition (vmPFC: *Z* = 4.45, *p*_perm_ < .001; PMC: *Z* = 4.09, *p*_perm_ < .001).

Pairwise comparisons revealed that both regions were sensitive to global context, reflected by significantly higher pattern similarity in the same context versus different context condition (vmPFC: Z = −4.49, *p*^perm^ < .001; PMC: Z = −3.88, *p*_perm_ < .001). Neither region was sensitive to local context, as reflected by no differences in pattern similarity between commercials from the same context versus similar context condition (vmPFC: *Z* = −1.95, *p*_perm_ = .056; PMC: *Z* = −1.43, *p*_perm_ = .150).

Turning to regions within the medial temporal lobe, contrary to our hypotheses, within the hippocampus, we observed no significant overall effect of condition (*Z* = 1.28, *p*_perm_ = .397). In medial temporal lobe cortical regions, the PARAHIPPOCAMPAL CORTEX showed a significant overall effect of condition (*Z* = 4.10, *p*_perm_ < .001), and pairwise comparisons reflected sensitivity to both global (*Z* = −4.15, *p*_perm_ < .001) and local context (*Z* = −2.72, *p*_perm_ = .011), with significantly higher pattern similarity for the same-context condition than the different- and similar-context conditions, respectively. In the perirhinal cortex, we observed a significant overall effect of condition (*Z* = 2.98, *p*_perm_ = .007), and pairwise comparisons showed sensitivity to global (*Z* = −3.08, *p*_perm_ = .008), but not local context (*Z* = −1.42, *p*_perm_ = .311); as in the cases of the other regions which were sensitive to global context, pattern similarity in the perirhinal cortex was significantly higher in the same-context condition than in the different-context condition.

Finally, we examined auditory cortex as a sensory control region and found no significant overall effect of condition (*Z* = 1.80, *p*_perm_ = .163). Pairwise comparisons in mean pattern similarity between conditions within each ROI are illustrated in Figure 2.

**Figure 2.**
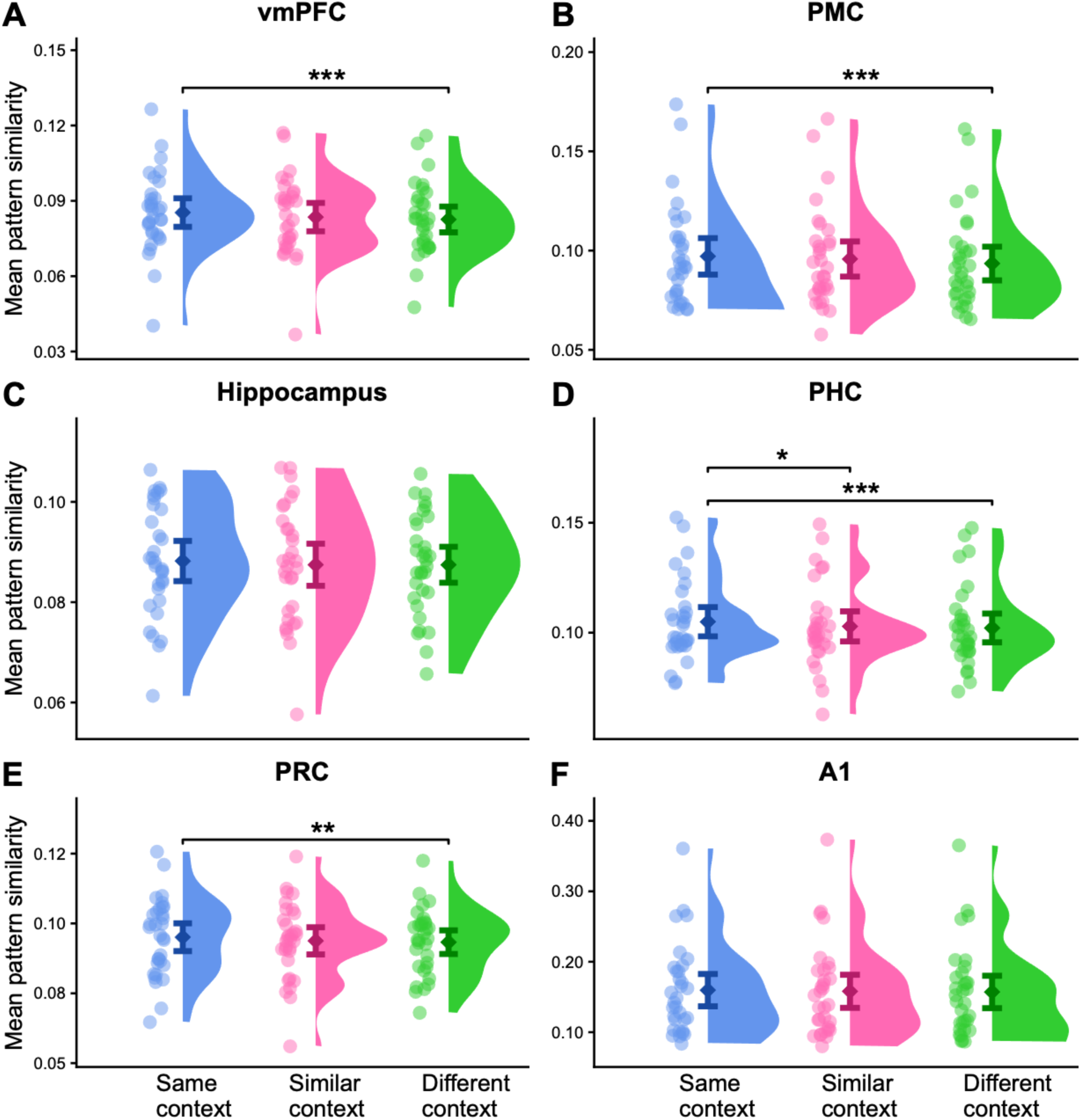
Pattern similarity. Raincloud plots show participant-level mean pattern similarity for commercial segments in the three conditions of interest: same context (same show, same episode), similar context (same show, different episode), and different context (different show). The ventromedial prefrontal cortex (vmPFC), posterior medial cortex (PMC), parahippocampal cortex (PHC), and perirhinal cortex (PRC) were sensitive to global context (same context significantly greater than different context, all *p*_perm_s ≤ .008). The parahippocampal cortex was additionally sensitive to local context (same context significantly greater than similar context, *p*_perm_ = .011). Data points represent individual participants. Dark diamonds represent means, and bars represent 95% confidence intervals. Raw correlation values are shown for visualization, but all statistical analyses were conducted on z-scored data. Statistical significance was assessed using permutation tests with 10,000 permutations. Significant comparisons are denoted with asterisks (* *p*_perm_ < .05; ** *p*_perm_ < .01; *** *p*_perm_ < .001).

#### Relationship between pattern similarity and source memory

To examine whether pattern similarity was related to subsequent source memory, we fit linear mixed-effects models including source memory accuracy, condition, and their interaction as predictors of pattern similarity. Consistent with our a priori hypothesis that the hippocampus would support memory for contextual associations, our analysis revealed a significant interaction between source memory accuracy and condition (*F*[2, 90] = 4.36, *p* = .016). Follow-up analyses of the slopes showed a positive association between source memory accuracy and hippocampal pattern similarity in the same-context condition (*t*[90] = 3.23, *p* = .002). The slope in the same-context condition was significantly greater than in the similar-context condition (Holm-adjusted *p* = .031) and the different-context condition (Holm-adjusted *p* = .035); the slopes in the similar-context and different-context conditions did not differ (Holm-adjusted *p* = .880). Thus, the positive association between source memory accuracy and hippocampal pattern similarity was specific to the same-context condition, indicating that greater hippocampal pattern similarity was associated with better source memory when commercials came from the same episode of the same TV show (see **Figure 3**). The interaction between source memory accuracy and condition was not significant in the remaining ROIs (PHC: *F*[2, 90] = 2.25, *p* = .111; PRC: *F*[2, 90] = 0.76, *p* = .472; vmPFC: *F[*2, 90] = 0.27, *p* = .763; PMC: *F*[2, 90] = 0.88, *p* = .419; A1: *F*[2, 90] = 2.16, *p*_m_ = .121).

**Figure 3.**
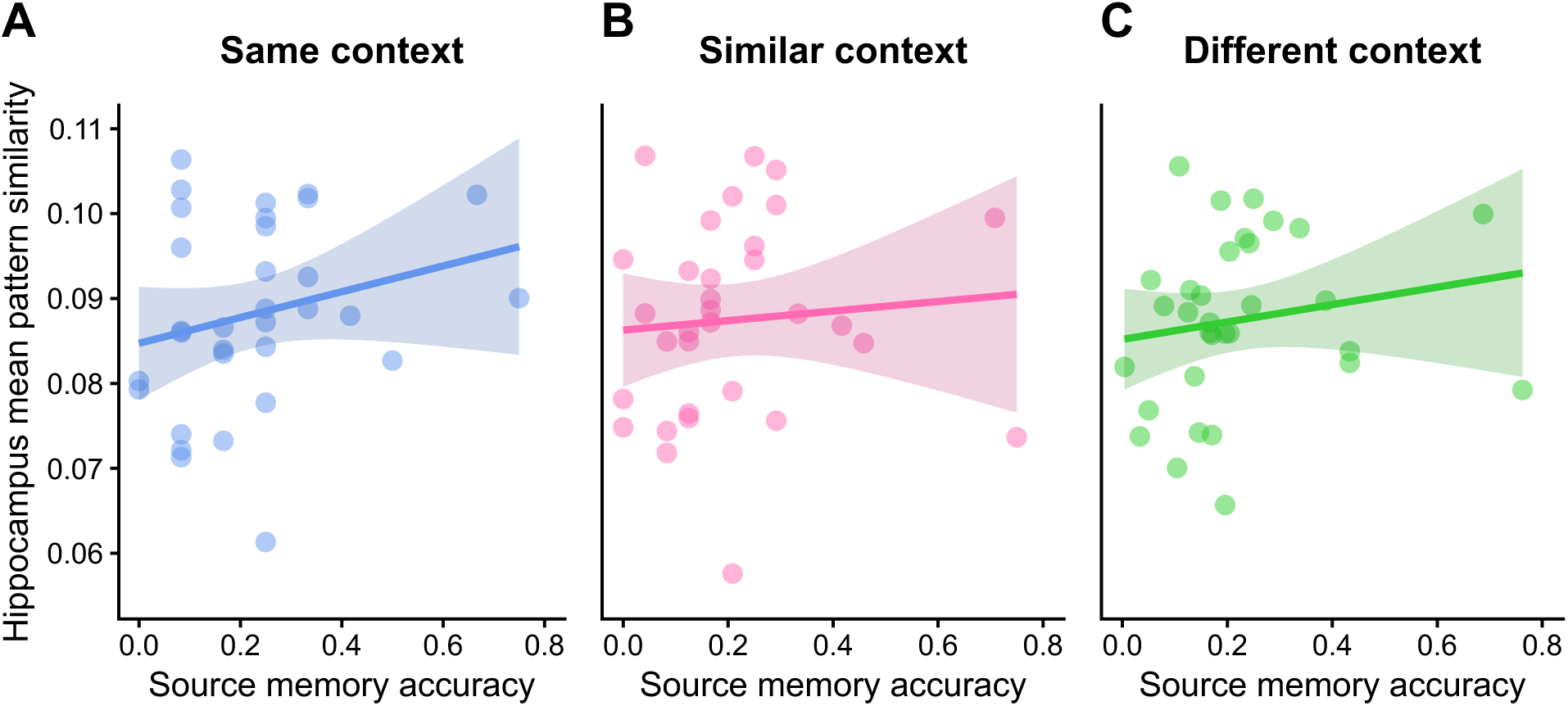
Relationship between hippocampal pattern similarity and source memory accuracy within each conceptual context category. Scatter plots show relationships between source memory accuracy and mean pattern similarity in the hippocampus for (A) same context, (B) similar context, and (C) different context. Solid lines represent the fitted linear regression for each condition, with shaded regions indicating 95% confidence intervals. Each point represents an individual participant. Source memory accuracy reflects the proportion of stimulus pairs for which participants correctly identified the source (TV show) for both commercials in the pair. Raw correlation values are plotted for visualization, but all statistical analyses were conducted on z-scored data.

However, we did observe a significant main effect of condition in the parahippocampal cortex (*F*[2, 90] = 3.57, *p* = .032), which aligns with our pattern similarity analyses and indicates that there was greater pattern similarity in the same-versus the similar-(Holm-adjusted *p* = .015) and same-versus different-context conditions (Holm-adjusted *p* = .003). Pattern similarity did not differ between the similar- and different-context conditions (Holm-adjusted *p* = .511).

In the remaining ROIs, we observed no main effects of either source memory accuracy (vmPFC, *F*[1, 90] = 0.29, *p* = .591; PMC, *F*[1, 90] = 0.05, *p* = .828; PRC, *F*[1, 90] = 1.59, *p* = .210; A1, *F*[1, 90] = 3.42, *p* = .068) or condition (vmPFC, *F*[2, 90] = 1.04, *p* = .359; PMC, *F*[2, 90] = 1.59, *p* = .209; PRC, *F*[2, 90] = 0.17, *p* = .846; A1, *F*[2, 90] = 2.40, *p* = .097).

#### Repetition suppression analyses

To examine whether repeated exposure to segments from the same commercial modulated neural representations without considering the influence of TV show context, we investigated conceptual repetition suppression effects in the commercial pre-exposure phase. This enabled us to explore how repetitions of commercial information across segments changed representations of the commercial. That is, for each commercial, three unique segments were seen during the pre-exposure phase. Importantly, individual segments were viewed just one time and no TV show context was present for any commercial. Paralleling classical repetition suppression effects, we reasoned that if, as someone saw additional segments from the same commercial, we detected a reduction in activity, that this would be evidence for sensitivity to shared conceptual information (Grill-Spector et al., 2006; Henson & Rugg, 2003).

Indeed, parahippocampal cortex showed evidence of this conceptual repetition suppression, with greater responses for segment 1 than segment 3 (FDR-corrected *p*_perm_ = .003), indicating this region was sensitive to repeated conceptual information. We did not observe significant effects in any other region (A1, *p*_perm_ = .900; hippocampus, *p*_perm_ = .363; PMC, *p*_perm_ = .363; PRC, *p*_perm_ = .592; vmPFC, *p*_perm_ = .363; all *p-*values FDR-corrected). Repetition suppression results for each ROI are illustrated in **Figure 4**.

**Figure 4.**
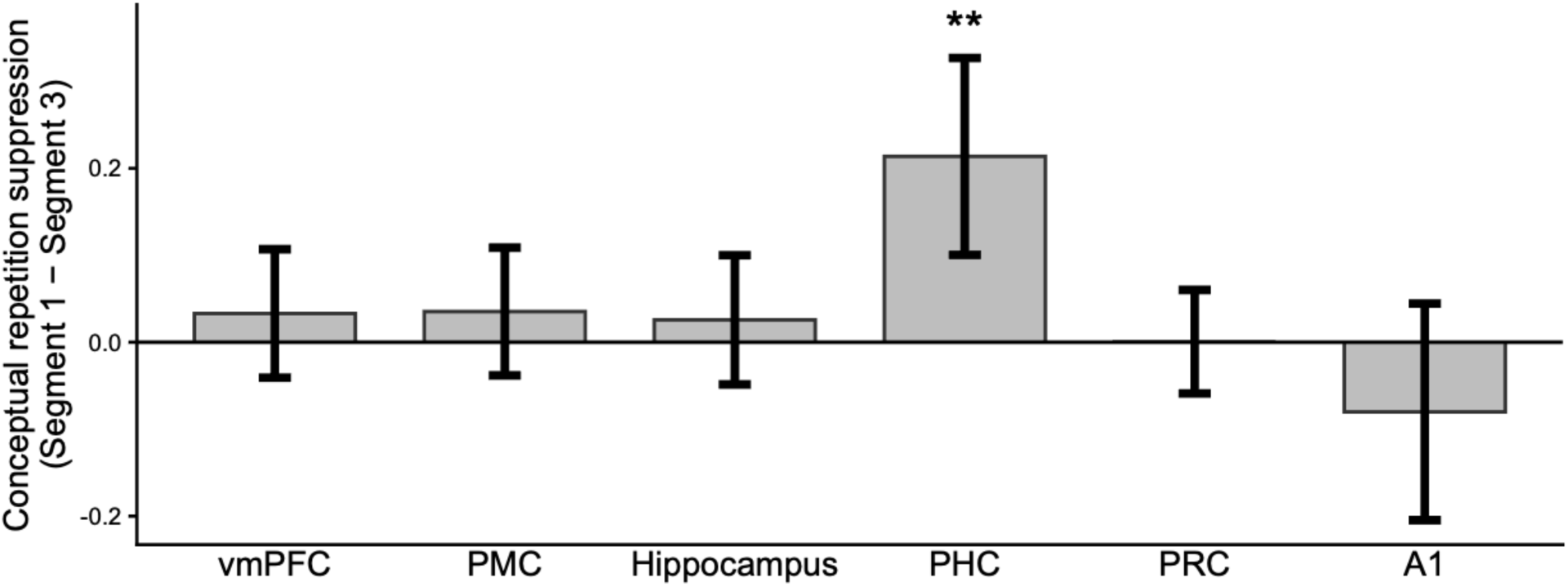
Repetition suppression. Conceptual repetition suppression, calculated as the difference in mean activation between segment 1 and segment 3 at the first presentation (Segment 1 − Segment 3) for each ROI. The parahippocampal cortex (PHC) showed a significant conceptual repetition suppression effect (Segment 1 > Segment 3). Bars represent means and error bars represent 95% confidence intervals. Significant effects are denoted with asterisks (** *p*_perm_ < .01).

## Discussion

While prior work has shown that encoding contexts both shape what gets remembered later (Hockley, 2008; Smith, 1979; Smith & Manzano, 2010; Tulving & Thomson, 1973) and are spontaneously reactivated at the time of retrieval (Bornstein & Norman, 2017; Bramão et al., 2017; Broitman & Kahana, 2026; Essoe et al., 2022; Lohnas et al., 2023; Manning et al., 2011; Masís-Obando et al., 2026) an understudied question is how events that share similar, but non-overlapping contextual associations, are represented relative to one another and how this may differ from representations for items from distinct contexts. To address this question in a way that mimics real-world experiences, we first exposed participants to commercials (events), devoid of any television show context. We next had participants watch these same commercials embedded within TV show episodes (contexts) while varying the degree of contextual overlap (same context: same TV show, same episode; similar context: same TV show, different episode; different context: different TV shows). Finally, to enable investigations of how our contextual manipulation changed representations, we once again showed commercials but without presenting any contextual information. At neither commercial viewing phase was any explicit memory retrieval demand imposed. When comparing changes in distributed activity profiles between the two commercial viewing experiences, we found evidence that both local context (which episode of the TV show) and global context (which TV show, regardless of episode) were represented.

In parahippocampal cortex, we found representations of both local context (same context > similar context) and global context (similar context > different context). In addition, when only considering the pre-TV exposure commercial viewings, parahippocampal cortex was sensitive to increasing levels of conceptual information about the commercials. In contrast to the parahippocampal cortex, the ventromedial prefrontal cortex and posterior medial cortex were sensitive to global, but not local, context. Notably, all of these regions have been suggested to play a role in representing context generally (Barnett et al., 2021; Eichenbaum, 2017a; Epstein & Baker, 2019; Hannula et al., 2013; Ritchey & Cooper, 2020; Turk-Browne et al., 2012) as well as in constructing overarching situation models that link across experiences (Buckner & DiNicola, 2019; Ranganath & Ritchey, 2012; Zwaan & Radvansky, 1998). Consistent with this view, the ventromedial prefrontal cortex has been implicated in representing schemas and higher-order structures that organize individual experiences (Eichenbaum, 2017b; Preston & Eichenbaum, 2013); for example, encoding-related activity patterns in the ventromedial prefrontal cortex have been associated with the representation of and memory performance for higher-order context information in narrative memory tasks (Baldassano et al., 2018; Masís-Obando et al., 2022). Similarly, activity in the posterior medial cortex has been associated with representations of broad contextual and event structure, such as the representation of event boundaries during movie watching at roughly the same granular level as human-made, natural-language annotations of event transitions (Baldassano et al., 2017; Chen et al., 2017).

Surprisingly, we also found that regions canonically affiliated with item-specific representations such as PRC (Davachi, 2006; Diana et al., 2012; Hannula et al., 2013; Ranganath, 2010; Staresina et al., 2011) were also sensitive to global contexts such that pattern similarity was greater for same-context commercials than for different-context commercials. Although the PRC is traditionally associated with item-level representations (Diana et al., 2012; Ranganath & Ritchey, 2012), it has also been implicated in learning of associations, especially if they can be linked to a single associated item or concept or unitized (Haskins et al., 2008; Staresina & Davachi, 2008), as well as in configural processing (Barense et al., 2007; Ladyka-Wojcik et al., 2026; A. C. H. Lee et al., 2005; Martin & Barense, 2023). Thus, the increased similarity in the PRC for same-context commercials may reflect shared conceptual or semantic structure associated with TV show context, rather than a representation of the specific episode-level context.

Intriguingly, and also contrary to our hypothesis, we did not observe any effects of context overlap in the hippocampus. Because prior work suggests that the hippocampus may play an integral role in binding items to contexts (Chanales et al., 2017; Davachi, 2006; Diana et al., 2007; Eichenbaum et al., 2007; Libby et al., 2019), we expected activity patterns in this region to be sensitive to local context. We reasoned that this lack of context representation in our data may be due to a lack of memory demand in the experimental task during the commercial viewing runs. To test this, we ran follow-up analyses examining the relationship between pattern similarity and source memory accuracy. This allowed us to determine whether hippocampal representations of context could be revealed when these representations were used in the service of supporting memory for contextual information. We found that greater pattern similarity was associated with better source memory for commercials from the same context condition (i.e., when both TV show and episode contexts were shared). This pattern suggests that the hippocampus may indeed play a role in local context representations, but only when memory for contextual information is preserved.

In sum, our findings provide evidence that neural representations of event contexts reflect the degree of contextual similarity between events, even when no explicit memory demand is present. Our approach allowed us to instantiate contextual associations between events within a continuous stream of experience and then measure the impact of these shared contexts on event representations measured at a separate point in time. We found that even when there was no explicit demand to retrieve contextual associations, item representations changed based on contextual affiliation. This suggests that context representations are carried alongside item representations and are an inherent part of item processing.

## Methods

### Participants

Thirty-four participants were recruited from the Columbia undergraduate and New York City community. They were compensated $60 for their time. This study was approved by the institutional review board at Columbia University. Two participants were excluded due to scanner malfunction. The results below reflect data from the remaining thirty-two participants (Mean age = 25.3 years, SD = 4.64 years, N_female_ = 20).

### Materials

Stimuli consisted of clips from one of six TV shows (*Corner Gas, Fillmore!, Pan Am, Popular, The Riches, The Tribe*). We selected these shows based on availability and likelihood that our participants would not be familiar with them. A researcher (HDZ) selected two episodes from each show where segments of approximately 90 seconds could be extracted. The goal was to ensure that each segment contained a relatively large scene boundary or shift; one commercial break was inserted here. A second commercial break was placed between episodes from different TV shows, as described below. In addition to the TV clips, 24 commercials were selected. To prioritize commercials that participants were unlikely to have seen, we searched for local commercials that were readily available from YouTube. Due to copyright, stimuli are not shared publicly but can be requested.

To create sequences where TV clips alternated with commercials, we then spliced together commercials and TV clips. The structure of this was such that participants saw clips from two different episodes from two different TV shows in each run (see ***Figure 1*** ‘TV watching phase’). The same two TV shows and the same commercials were always shown together within a run, although the order of the runs was randomized across participants. TV show pairings were as follows: Corner Gas/The Riches, Fillmore/The Tribe, Pan Am/Popular. Selection of TV show pairings and commercials was done randomly at the discretion of the researcher.

Within each run, clips from the two TV shows alternated such that the first episode of TV show A was presented, followed by the first episode of TV show B, then the second episode of TV show A, followed by the second episode of TV show B, and so on. Each episode clip included one commercial break, which was randomly placed by the experimenter with the goal of positioning it at a moment that would not disrupt understanding of the ongoing TV clip narrative (e.g., commercials were not placed in the middle of a conversation between two characters) although placement of this commercial was unlikely to have been at a major scene shift when you might have naturally expected there to be a commercial break. The intention was for these within-show commercial breaks to be weakly predicted by changes in the show’s content. Additionally, a commercial break was also placed between TV shows. This between-show commercial break was intended to be at a more natural or predictable location for a commercial break (although, as verified by participant recall reports, the success of the researcher’s intentions varied by clip/commercial placement). Since all participants saw the same TV clips with the same commercial placement, we do not feel that this limitation substantially contributed to the observed pattern of results.

In addition, we probed participants about the “fit” of the commercial with the previous TV clip (see *Procedure* for more details). A prompt to make this judgment appeared on the screen during each commercial, and the participant responded via button press on an MRI-compatible button box. For each TV show run, we created ten versions in which the timing of each commercial probe varied; for each participant, we randomly drew one of the ten options for each run. We ensured that there was always a 10-second buffer from the beginning and a 5-second buffer from the end of the commercial before the fit questions were delivered, to allow the participant to fully view the commercial before making their fit decision. In addition, across participants, we varied which response option appeared on the left or right of the screen (i.e., sometimes “bad” appeared on the left, and other times on the right). Left/right button placement did not vary within a participant.

### Procedure

Prior to entering the MRI scanner, participants were introduced to the task by a set of instruction slides. The experimenter told participants that they would watch video clips and commercials and make judgments about the “fit” of the commercials with those shows.

Participants were instructed to imagine that they had been hired by a marketing research firm to judge whether the commercial was in a “good timeslot” or a “bad timeslot”. Participants were instructed that they could use the content of the TV shows and commercials, the timing of the commercial within a show clip, or some combination of those features to make this judgment. However, the experimenter stressed that they should use a consistent heuristic to make these fit judgments throughout the task. Participants were also instructed that, after viewing the TV episode clips and commercials, they would be asked to recall aloud everything they could remember.

Participants then completed a final metal screening check and entered the MRI room. In addition to the standard MRI setup that included a Psychology Software Tools 4 button response pad and safety squeeze ball, participants were also fitted with Optoacoustics OptoActive II noise-canceling headphones or Avotec Silent Scan 3300 noise-canceling earbuds that delivered the audio from the TV shows and commercials (various options were used across participants due to recurrent technical issues with audio presentation). We checked that participants could hear the audio well and in both ears before showing them any of the experimental stimuli. Of note, this sometimes required several attempts; for some participants, audio quality remained poor. To enable recording of free recall in the scanner, we used an Optoacoustics OptoActive II system microphone that was mounted to the outside of the head coil (see Soares et al., 2024 for similar setup and procedures). A rear-facing mirror was fitted on top of the head coil to ensure participants could see the TV shows projected onto the screen.

Once set up inside the MRI scanner, participants were asked to view trailers of each TV show, read a synopsis, and “meet” key characters (by seeing pictures of them paired with names) from each of the six TV shows they would later be seeing clips from. This section was self-paced and took place during the acquisition of the structural (T1 and T2) MRI scans.

#### Commercial pre-exposure phase

After the TV show introductions and structural scans were completed, participants were told they would complete a task to help calibrate the scanner. They were told they would view short clips from the commercials they would see later, and that their job was to press a button if they detected that any clips shown back-to-back were from the same commercial.

Participants viewed 72 5-second clips from each commercial in this phase. The order of these clips was uniquely randomized for each participant using the ‘random.permutation’ function from the NumPy library. Since this was fully randomized, seeing two clips from the same commercial occurred infrequently and may not have occurred for each participant. This cover story was meant to keep participants engaged and watching the clips. Clips were separated with a 5-second inter-stimulus interval.

#### TV watching and recall phases

Following the commercial pre-exposure phase, participants were reminded of the commercial fit judgment instructions and then proceeded to the TV watching and recall phase. In each run, participants watched clips from four TV episodes (two per show) and saw 8 commercials. TV shows and commercials were unique to each run. At some point during the commercial, a black box appeared at the bottom of the screen with the response options “good timeslot” and “bad timeslot” (the timing of this probe within the commercial varied across participants and the side of the screen for each response option was counterbalanced; see description above). Participants were instructed to respond, via keypress, about their judgment of the fit of the commercial with the TV clip that had been seen immediately before the commercial. We hoped this task would both engage participants (all participants reported that the stimuli were quite interesting, and we had no issues with engagement) and help participants make an explicit link between the TV show and the commercial.

Following the final commercial, a black screen appeared that said “Waiting for recall phase to start…”. This screen remained up for 22-29 seconds. This ensured that the TV watching phase took up 10 minutes and 30 seconds for each run (i.e., since there was slight timing variability between the TV clips from different shows, we wanted to match the full length of this TV watching phase). This phase is not described further here since data are not reported. These TV watching and recall procedures were repeated, with breaks between runs as necessary, for three runs.

#### Commercial post-exposure phase

This phase was identical to the pre-exposure phase except that the video clips were randomized in a different order than in the pre-exposure phase (however, all clip identities remained the same in both phases).

#### Post-experiment memory phase

Outside of the scanner, participants completed a memory test for the commercial-TV show pairings. They were presented with one still frame from each of the 24 commercials and indicated, via button press, which of the six TV shows they remembered the commercial having been paired with during the TV-watching run. No feedback about the accuracy of these responses was given.

### Behavioral data analysis

Source memory performance was assessed using the post-experiment memory test. For each commercial, participants’ responses were scored as correct or incorrect based on whether they correctly identified the commercial’s source (i.e., which TV show it was paired with during the TV-watching run). Overall memory accuracy was calculated as the proportion of commercials whose affiliated TV show source was correctly identified across the memory test.

We then assessed source memory at the level of the predefined commercial pairs used in the pattern-similarity analyses. For each pair, successful source memory was classified as when both commercials were correctly identified.

### fMRI data analysis

Data were collected on a 3T Siemens Magnetom Prisma MRI scanner. Anatomical images were acquired using a T1-weighted MPRAGE sequence. Images were acquired with a 1mm isotropic resolution (TR = 2300 ms; TE = 2.98 ms; FOV = 256 × 249 × 176 mm; flip angle = 9°) and in the sagittal orientation with anterior-to-posterior (A-P) phase encoding direction. Functional images were acquired with 2.0-mm isotropic voxels in an oblique orientation with anterior-to-posterior (A-P) phase encoding direction. Acquisition parameters were TR = 1500 ms, TE = 28 ms, and flip angle = 65°. A total of 494 volumes were acquired across 69 interleaved slices.

Results included in this manuscript come from preprocessing performed using fMRIPrep 21.0.1 (Esteban, Blair, et al., 2019; Esteban, Markiewicz, et al., 2019), which is based on Nipype 1.6.1 (K. Gorgolewski et al., 2011; K. J. Gorgolewski et al., 2018). We have minimally edited the fMRIPrep-generated text for readability.

#### Anatomical data preprocessing

The T1-weighted (T1w) image was corrected for intensity non-uniformity (INU) with N4BiasFieldCorrection (Tustison et al., 2010), distributed with ANTs 2.3.3 (Avants et al., 2008), and used as the T1w reference throughout the workflow. The T1w reference was then skull-stripped with a Nipype implementation of the antsBrainExtraction.sh workflow (from ANTs), using OASIS30ANTs as the target template. Brain tissue segmentation of cerebrospinal fluid (CSF), white matter (WM), and gray matter (GM) was performed on the brain-extracted T1w using FAST (FSL 6.0.5.1; Zhang et al., 2001). Brain surfaces were reconstructed using recon-all (FreeSurfer 6.0.1; Dale et al., 1999), and the brain mask estimated previously was refined with a custom variation of the method to reconcile ANTs-derived and FreeSurfer-derived segmentations of the cortical gray matter of Mindboggle (Klein et al., 2017). Volume-based spatial normalization to one standard space (MNI152) was performed through nonlinear registration with antsRegistration (ANTs 2.3.3), using brain-extracted versions of both the T1w reference and the T1w template. The following template was selected for spatial normalization: ICBM 152 Nonlinear Asymmetrical template version 2009c (Fonov et al., 2009).

#### Functional data preprocessing

For each subject, the following preprocessing was performed on the commercial pre-and post-exposure runs: first, a reference volume and its skull-stripped version were generated using a custom methodology of fMRIPrep. Head-motion parameters with respect to the BOLD reference (transformation matrices and six corresponding rotation and translation parameters) were estimated before any spatiotemporal filtering using mcflirt (FSL 6.0.5.1; Jenkinson et al., 2002). The BOLD time-series (including slice-timing correction when applied) were resampled onto their original, native space by applying the transforms to correct for head motion. The BOLD reference was then co-registered to the T1w reference using bbregister (FreeSurfer), which implements boundary-based registration (Greve & Fischl, 2009). Co-registration was configured with six degrees of freedom. Several confounding time-series were calculated based on the preprocessed BOLD: framewise displacement (FD), DVARS, and three region-wise global signals. FD was computed using two formulations following Power (absolute sum of relative motions; Power et al., 2014) and Jenkinson (relative root mean square displacement between affines; Jenkinson et al., 2002). FD and DVARS were calculated for each functional run, both using their implementations in Nipype (following the definitions by Power et al., 2014). The three global signals were extracted within the CSF, the WM, and the whole-brain masks. Additionally, a set of physiological regressors was extracted to allow for component-based noise correction (CompCor; Behzadi et al., 2007). Principal components were estimated after high-pass filtering the preprocessed BOLD time-series (using a discrete cosine filter with 128s cut-off) for the two CompCor variants: temporal (tCompCor) and anatomical (aCompCor). tCompCor components were then calculated from the top 2% variable voxels within the brain mask. For aCompCor, three probabilistic masks (CSF, WM and combined CSF+WM) were generated in anatomical space. The implementation differs from that of Behzadi et al. (2007) in that instead of eroding the masks by 2 pixels in BOLD space, the aCompCor masks were subtracted from a mask of pixels that likely contain a volume fraction of GM. This mask is obtained by dilating a GM mask extracted from the FreeSurfer’s aseg segmentation, and it ensures components are not extracted from voxels containing a minimal fraction of GM. Finally, these masks were resampled into BOLD space and binarized by thresholding at 0.99 (as in the original implementation). Components were also calculated separately within the WM and CSF masks. For each CompCor decomposition, the k components with the largest singular values were retained, such that the retained components’ time series are sufficient to explain 50 percent of variance across the nuisance mask (CSF, WM, combined, or temporal). The remaining components were dropped from consideration. The head-motion estimates calculated in the correction step were also placed within the corresponding confounds file. The confound time series derived from head motion estimates and global signals were expanded with the inclusion of temporal derivatives and quadratic terms for each (Satterthwaite et al., 2013). Frames that exceeded a threshold of 0.5 mm FD or 1.5 standardized DVARS were annotated as motion outliers. The BOLD time-series were resampled into standard space, generating a preprocessed BOLD run in MNI space. First, a reference volume and its skull-stripped version were generated using a custom methodology of fMRIPrep. The BOLD time-series were resampled onto the following surfaces (FreeSurfer reconstruction nomenclature): fsaverage6. All resamplings were performed with a single interpolation step by composing all the pertinent transformations (i.e., head-motion transform matrices, susceptibility distortion correction when available, and co-registrations to anatomical and output spaces). Gridded (volumetric) resamplings were performed using antsApplyTransforms (ANTs), configured with Lanczos interpolation to minimize the smoothing effects of other kernels (Lanczos, 1964). Non-gridded (surface) resamplings were performed using mri_vol2surf (FreeSurfer).

#### First-level fMRI specification

First-level models were specified using SPM12 (v7771; Statistical Parametric Mapping; Welcome Centre for Human Neuroimaging, University College London). Preprocessed BOLD images generated with fMRIPrep and normalized to MNI space were entered into subject-level models. Experimental conditions were specified using subject- and condition-specific timing files containing the onset and duration information. Each condition was modeled by convolving its event timings with the canonical hemodynamic response function (HRF) without temporal or dispersion derivatives. To account for variance associated with head motion and non-neuronal noise, subject-specific nuisance regressors derived from the fMRIPrep preprocessing pipeline were included in the first-level model. These included six rigid-body head-motion parameters (three translational parameters: x, y, and z; and three rotational parameters: x, y, and z) and their first temporal derivatives. Framewise displacement was also included as an additional nuisance regressor to account for motion-related signal variance. Additionally, six anatomical component-based noise correction (aCompCor) components (aCompCor_00–aCompCor_05) were included as nuisance regressors. A 128-s high-pass filter was applied to remove low-frequency signal fluctuations, and temporal autocorrelation was modeled using an AR(1) process. No global signal normalization was applied.

The commercial pre-exposure and post-exposure phases were modeled separately to obtain independent, run-specific beta estimates. Segment-level patterns were estimated with a least-squares-separate (LSS) approach (Mumford et al., 2014). For each commercial, three GLMs were fit, one per segment. Each GLM included a 5-second boxcar regressor for the segment of interest, a regressor for the remaining two segments of the same commercial, and a regressor for all segments of the other 23 commercials. With 24 commercials and each commercial potentially contributing one of three segments as the segment of interest, this procedure yielded 72 first-level models per participant for each pre-and post-exposure run.

#### Regions of interest

We determined six a priori regions of interest (ROIs; see **Figure 5**) that were used in subsequent analyses. The hippocampus, parahippocampal cortex, and perirhinal cortex were defined using automated segmentation from the Automatic Segmentation of Hippocampal Subfields (ASHS; version 1.0.0, with the Princeton atlas) framework. ASHS returns hippocampal subfields as separate labels. However, because our scans were not high-resolution, we were unable to reliably isolate functional signal to individual subfields. Therefore, we combined CA1, CA2/3, dentate gyrus, and subiculum subfield labels to create a single, subject-specific hippocampal mask for each hemisphere separately; left and right hemispheres were then merged for analyses. The ventromedial prefrontal cortex (vmPFC) ROI was derived from the MNI-BIC vmPFC atlas (https://www.bic.mni.mcgill.ca/ServicesAtlases/VmPFC). The posterior medial cortex (PMC) ROI was obtained from the Baldassano laboratory (personal communication). Finally, the primary auditory cortex (A1) was included as a control ROI and was defined using the extended Human Connectome Project multimodal parcellation atlas (HCPex; Huang et al., 2022). Auditory cortex was selected as a control region because it supports relatively early-stage auditory sensory processing and was not expected to be specifically involved in the higher-order contextual and mnemonic processes targeted by our hypotheses.

**Figure 5.**
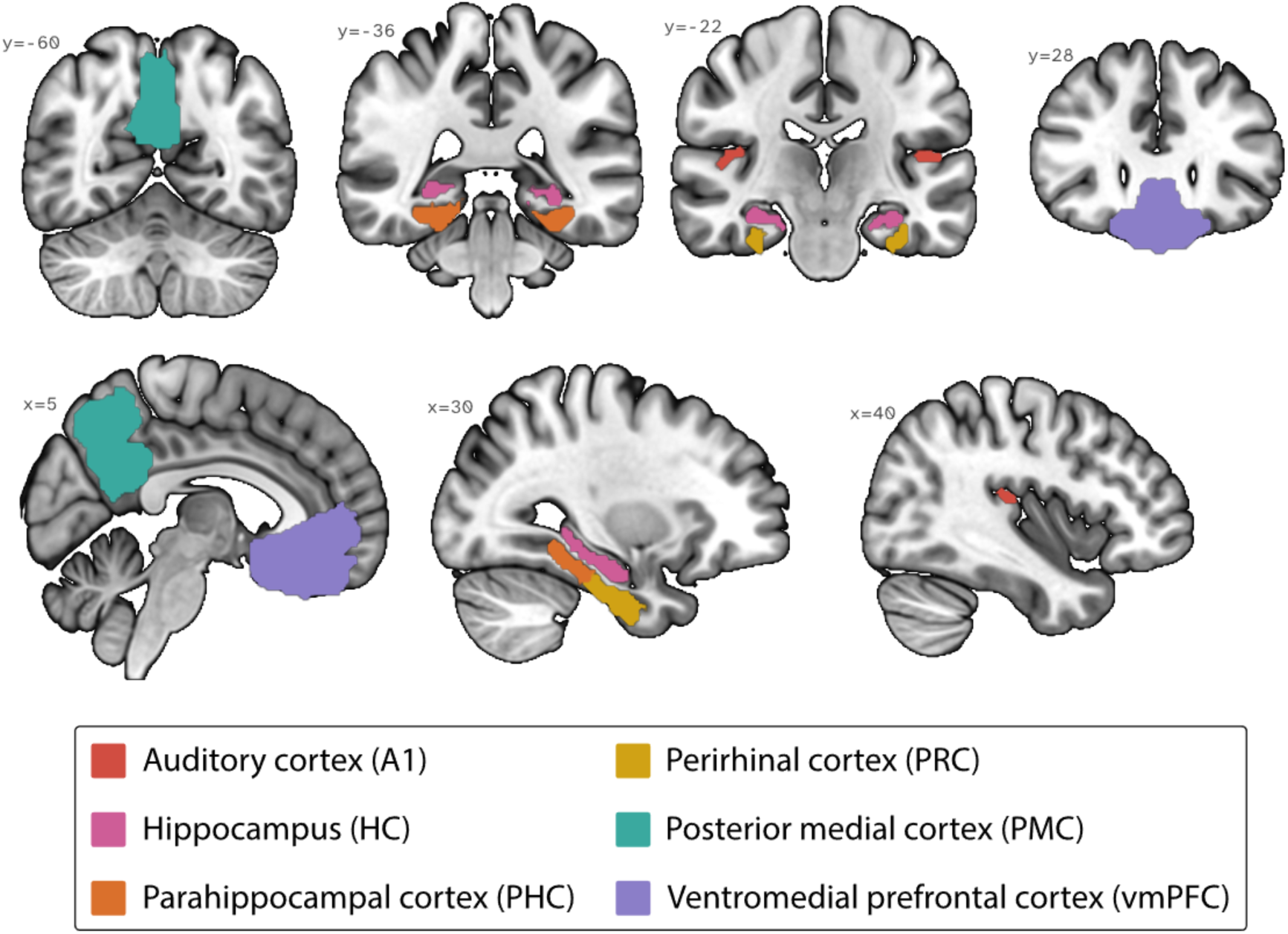
Visualization of regions of interest (ROIs) in coronal and sagittal planes. Regions of interest included the hippocampus (HC), parahippocampal cortex (PHC), perirhinal cortex (PRC), posterior medial cortex (PMC), and ventromedial prefrontal cortex (vmPFC). The primary auditory cortex (A1) was included as a control region. ROIs are overlaid on the MNI brain using MRIcroGL (https://www.nitrc.org/projects/mricrogl/; Brett et al., 2001). Slices are displayed in neurological convention and labeled with corresponding slice coordinates.

For each ROI, left- and right-hemisphere masks were generated separately and used to extract beta estimates. Because we had no a priori hypotheses regarding hemispheric differences, data were collapsed across hemispheres within each ROI for all reported analyses. All ROI masks were binarized and resampled to the functional image space prior to extraction of beta estimates.

#### Beta-weight extraction and pattern similarity analysis

For each participant and ROI, beta estimates were extracted for each commercial segment of the 24 commercials from the corresponding first-level models, separately for the pre-exposure and post-exposure phases. This resulted in a voxel-by-commercial segment matrix for each participant, ROI, and phase (i.e., pre/post). To quantify changes in multivoxel response patterns from pre-to post-exposure, beta estimates were z-scored across commercials for each voxel for the pre- and post-exposure runs. We then calculated the difference between the pre- and post-exposure patterns (POST − PRE). Voxels containing missing values were excluded (this occurred infrequently). Pearson correlations were then calculated between the resulting POST– PRE difference scores for every pair of commercials, yielding a 24 x 24 correlation matrix for each participant and ROI. Thus, each cell of the resulting matrix represented the similarity between the changes in multivoxel response patterns associated with commercial pairs. These correlation matrices were used for the pattern similarity analyses.

Pattern similarity values were analyzed separately within each ROI using R (version 4.5.2). Each pair of commercials was classified according to its relationship to the TV shows they were paired with during the TV watching run: (1) same TV show and same episode (same context), (2) same TV show and different episode (similar context), (3) different TV show (different context). To test whether pattern similarity differed across these conditions, we conducted within-subject permutation tests using the independence_test() function from the coin R package (version 4.5.2). Of note, we used permutation-based inference because the analyses were based on the trial-level pattern-similarity estimates and the number of observations contributing to each condition differed substantially. The test evaluated the null hypothesis of no association between condition and pattern similarity while accounting for the repeated-measures structure of the data by restricting permutations within participants. Overall tests were conducted separately for each ROI. For ROIs showing a significant overall effect, post hoc pairwise permutation tests were conducted for all three pairwise comparisons between conditions. Pairwise p-values were adjusted for multiple comparisons using the Holm correction. Each permutation test was approximated using 10,000 resamples, and the reported test statistic was the standardized linear statistic derived from the permutation distribution.

#### Pattern similarity and source memory analysis

The pattern similarity correlation estimates used in this analysis were the POST − PRE difference scores described above. Source memory performance was assessed using the post-scan memory test. For each participant and condition, correlation estimates were averaged across stimulus pairs to obtain a participant-level measure of mean neural pattern similarity, separately for each ROI. To examine the relationship between neural pattern similarity and source memory performance while accounting for the repeated observations contributed by each participant across conditions, linear mixed-effects (LME) models were fit separately for each ROI.

Mean neural pattern similarity was modeled as a function of source memory accuracy, condition, and their interaction, with participant entered as a random intercept. Our primary hypothesis concerned the interaction between source memory accuracy and condition, testing whether the relationship between neural pattern similarity and source memory differed across the three experimental conditions (i.e., same TV show and same episode, same TV show and different episode, different TV show). Significant main effects were further explored with pairwise comparisons between conditions to estimate marginal means, with Holm correction for multiple comparisons; significant interactions were examined with condition-specific slopes that were estimated using estimated marginal trends, and pairwise comparisons of slopes were conducted with Holm correction.

The LME models for each ROI therefore tested (1) whether neural pattern similarity was related to source memory accuracy, (2) whether mean neural pattern similarity differed across conditions, and (3) whether the relationship between source memory accuracy and neural pattern similarity differed across conditions.

#### Repetition suppression analysis

Repetition suppression refers to the reduction in neural response when the same stimulus is presented repeatedly, typically assessed by comparing the neural response to an initial presentation with the response to a subsequent presentation of that same exact stimulus (Grill-Spector et al., 2006; Henson & Rugg, 2003). In the present study, however, we did not operationalize repetition suppression in this way. Doing so would have required directly comparing neural responses to the same commercial segment across the pre- and post-exposure runs. Because these sessions were acquired in separate scanning runs and were separated by several intervening tasks (i.e., the 3 TV-watching runs and the free recall task), we could not rule out the possibility that the extended time delay between the 2 stimulus presentations or other factors associated with the intervening tasks would have influenced the changes in neural response.

Instead, we examined conceptual repetition suppression, which we operationalized as a reduction in neural responses across stimuli that share conceptual content rather than being identical. In other words, conceptual repetition suppression was operationalized as the difference in neural response between the first segment (seg-1) and the third segment (seg-3) of a commercial as presented during the initial pre-exposure run, regardless of their chronological position within the original commercial. This comparison was intended to capture reduced neural responding to a segment following exposure to conceptually related content within the same commercial. Positive values therefore indicated greater neural response to the initial segment than to the later conceptually related segment. For each ROI, these difference scores were tested against zero using one-tailed one-sample t-tests, with the directional hypothesis that neural responses would be reduced for the later conceptually related segment. For each ROI, these difference scores were tested against zero using one-tailed permutation tests based on 10,000 random sign permutations.

